# Super-resolved mass photometry reveals the dissociation pathways of individual protein complexes

**DOI:** 10.64898/2026.11.14.766467

**Authors:** Alexander R. O’Shea, Josh L. Bishop, Franziska L. Sendker, Jack Bardzil, Mahi Patel, Jack S. Peters, Max F. Hantke, Masahiro Suekawa, Tetsuo Torisu, Susumu Uchiyama, Georg K. A. Hochberg, Philipp Kukura, Justin L. P. Benesch

## Abstract

Mass photometry (MP) measures the mass of single unlabelled molecules as they adsorb to a glass surface, but conventional analysis captures each molecule only momentarily, as it binds, and neglects its subsequent fate. Here we correct for coverslip drift and use mass-dependent localisation precision to link successive events at one site to a single molecule and enables two workflows for studying protein assemblies. In tandem MP, matching each binding event with its outcomes yields a two-dimensional precursor-product spectrum that separates dissociation from unbinding, providing structural insights into the protein assemblies and the kinetics of subunit loss. By comparing citrate synthase homologues, we reveal distinct fingerprints and disassembly pathways among complexes of the same oligomeric state. In particle tracking, differential imaging allows for continuous recording of particle mass and position. By examining adeno-associated virus capsids, we recovered accurate masses, and reveals acid-induced disassembly and movement behaviour that depends on both viral and imaging surfaces. Our code is freely available and applies to data obtained on commercial instruments with minimal changes to experimental protocols, offering additional dimensions to MP analysis beyond the mass histogram to return time-resolved, single-molecule histories.

## Main

The function of proteins and their higher-order structures depends on their assembly, dynamics, and transitions between various states^1,2^. Structural biology methods such as cryo-electron microscopy, nuclear magnetic resonance spectroscopy, and X-ray crystallography can resolve composition with high precision^3–5^, but typically average over many molecules to provide an ensemble average and struggle to monitor molecular transitions. Single-molecule techniques can report on the underlying heterogeneity that ensemble measurements average away but typically require the molecule to be labelled with fluorophores or other tags^6,7^.

Mass photometry (MP) is a label-free in-solution optical mass measurement technique that has proven effective for measuring biomolecular stoichiometry^8,9^ and quantifying interactions^10^. In a typical MP experiment, a camera records the change in reflectivity of a glass cover-slip, as single molecules from solution bind to it. Specifically, biomolecules in solution scatter light as they adsorb onto the glass surface, and this interferes with the surface-reflected field, allowing individual molecules to be detected with a contrast signal proportional to mass^8^. This signal is quantified using ratiometric imaging: a “rolling window” of video frames is divided by a window preceding it to suppress the static background that dominates the detected signal^11^. Ratiometric videos reveal binding events as temporary features, with the magnitude of contrast increasing as a biomolecule’s signal lands in the running window and decreasing once it is present in both windows^12^. As a result, each individual molecule is effectively only measured once (with the maximum contrast extracted and converted to a mass); after the ratiometric window has passed, the signal of the bound particle is neglected, even though the particle continues to scatter for as long as it remains on the surface.

To mass-measure individual molecules for extended periods of time, a variant of MP was developed whereby molecules attached to a supported lipid bilayer could be monitored as they diffused in the bilayer plane^13,14^. This allows the continuous mass measurement of molecules by substituting a ‘rolling median’ background for ratiometric processing and converting lateral mobility across the bilayer into a diffusion coefficient orthogonal to mass. Recently, single molecule trapping extended the observation time of a single molecule diffusing on a bilayer from a few seconds to hours^15^, allowing molecular transitions to be monitored with exquisite resolution^16^. However, this approach also comes with limitations: 1) it requires the molecules to be tagged so they can be linked to the lipid bilayer; 2) generating supported lipid bilayers is cumbersome, and they are unstable on current commercial mass photometers, 3) only a handful of molecules can be measured at the same time as their lateral diffusion leads to crossing trajectories, and 4) the bilayer itself produces substantial additional imaging background, raising the detection limit to >100 kDa.

Here we introduce a novel approach for measuring the behaviour of individual molecules by means of MP. We reasoned that the fate of adsorbed particles is encoded in the raw video of the glass surface, despite a binding event is only transiently observed in ratiometric imaging and them also appearing absent from a rolling median image (if they do not move further than the diffraction limit of light). However, we noted that it is non-trivial to differentiate between successive events at seemingly the same position originating from the same molecule from coincidental events occurring to an unrelated molecule nearby. Lateral drift of the glass coverslip over the course of a measurement, as well as potential movements of the molecules themselves, compounds this ambiguity. To overcome these challenges, and allow extended measurements of individual proteins on glass, we present Python code to correct for glass drift over time, quantify localisation precision, and to accommodate movements of molecules on the glass. This work extends MP analysis to two new workflows: 1) tandem MP, where the binding, dissociation and unbinding of individual molecules can be linked, and 2) particle tracking, where the trajectories that molecules take on the coverslip can be quantified to reflect their surface chemistry.

Tandem MP operates through pair-matching an event to its successor at the same location using a localisation precision model, producing a two-dimensional precursor-product spectrum that separates dissociation from unbinding and resolves subunit-loss stoichiometry and kinetics. We validate tandem MP against the known assembly of β-galactosidase and apply it to a family of citrate synthases^17,18^, revealing that members of the family with the same stoichiometry distribution display differences in their disassembly pathways, and quantifying the kinetics of dissociation are impacted by the binding of substrate. In particle tracking, our software continuously monitors adsorbed particles using differential imaging with a dynamically updated background image, producing simultaneous orthogonal readouts of mass and position. In experiments on adeno-associated viruses (AAVs), we demonstrate that this approach allows movements on the surface, which can give rise to artefacts in ratiometric imaging, to be recovered into accurate measurements of mass. We also demonstrate that we can quantify the disassembly of AAVs, how it depends on pH, and show that different AAV serotypes move on the surface at different speeds, reflecting differing surface chemistry. Together, our advances in data analysis recover information that conventional analysis currently neglects, enabling the quantification of the dynamic behaviour of hundreds of unlabelled protein assemblies in parallel, in a simple MP experiment.

## Results

### Super-resolved MP allows the linking of multiple events at the single molecule level

MP measures the mass of single molecules^8,19^ and their position on the glass surface, with nanometre precision^19,20^. In MP convention, biomolecules binding *on* the surface, produce negative (dark) contrast, while biomolecules detaching *off* the surface produce contrast of the opposite sign, positive (bright) contrast. In principle, assuming each biomolecule (precursor) remains in the same diffraction-limited spot once bound, its *binding* event can be linked, or ‘paired’, to any subsequent dynamic transitions it undergoes (**Fig. 1a**). There are three potential transitions: *unbinding*, where the whole biomolecule leaves the surface, producing an event of the same contrast but opposite sign (**Fig. 1a, left**); *dissociation*, where the biomolecule sheds one or more (product) subunits, producing an unbinding event with a smaller contrast (**Fig. 1a, middle**); and *movement*^21^, where the biomolecule relocates to a neighbouring diffraction-limited position, producing a characteristic part-bright, part-dark signal representing unbinding at one position followed by rapid re-binding at another (**Fig. 1a, right**). We aimed to develop a means of pairing these events, so that faithful histories of biomolecules could be traced.

**Fig. 1.**
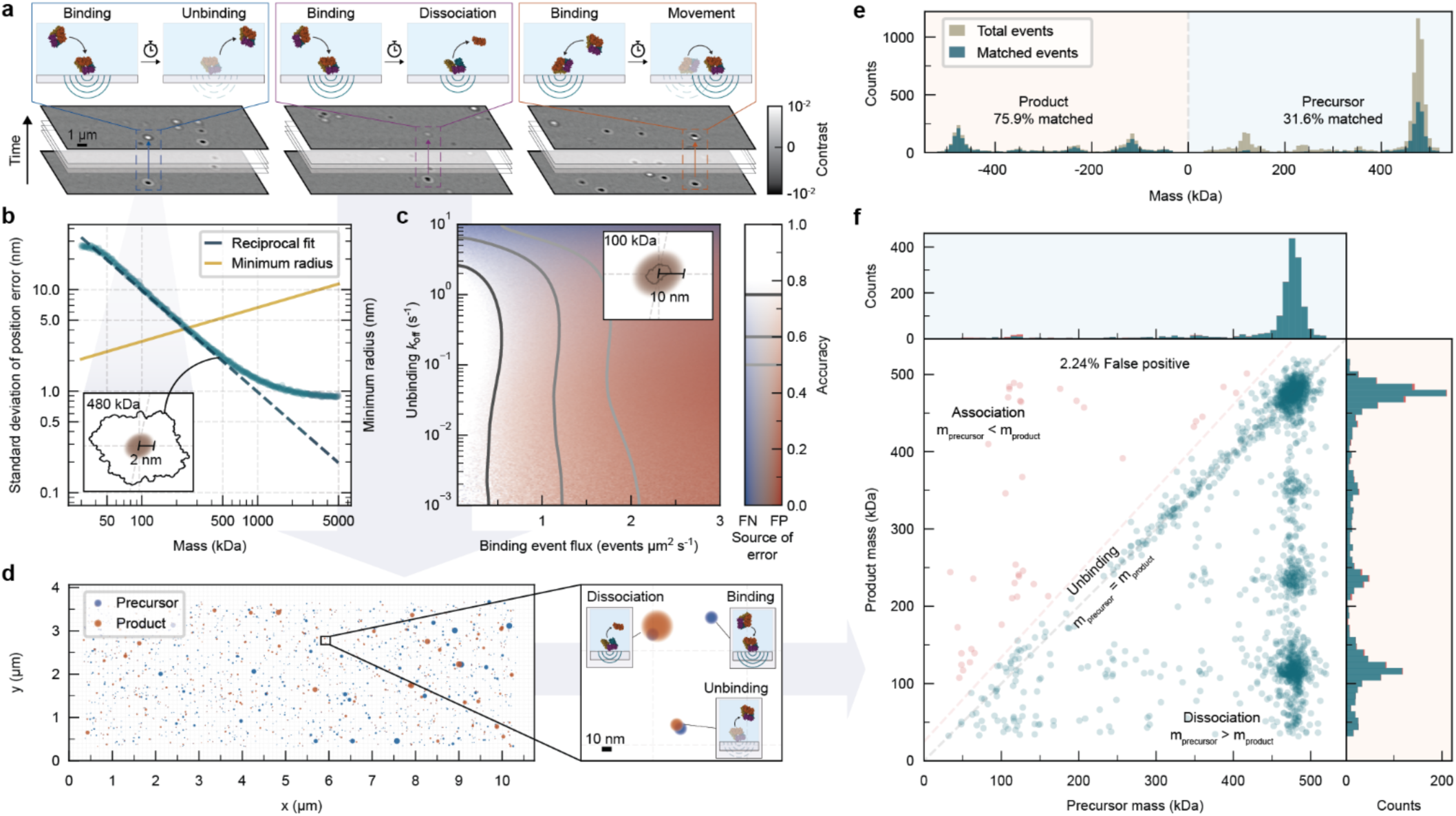
Principle of pair-matching: linking binding events to their outcomes using a localisation precision model. **a**, Schematic illustrating three different outcomes that can occur once a biomolecule binds to the glass surface in a mass photometry movie: left, intact unbinding; middle, dissociation; right, movement. Dashed boxes track the same position across frames taken from a β-galactosidase (*E. coli*) measurement. **b**, Mass dependent localisation precision obtained by fitting a mathematical PSFs to simulated PSFs of varying mass on experimental backgrounds (obtained using an integration time of 100ms^23^). Each point represents the standard deviation of the position error distribution formed by 1,000 repeats. A reciprocal fit consistent with shot-noise-limited scaling is shown by the teal dashed line. Theoretical minimum radius of a protein sphere of given mass is shown by the gold line. Inset: to-scale representation of the outline of a β-galactosidase tetramer complex (PDB ID: 4V40, mass ∼464 kDa) and the localisation precision of a 480 kDa event. **c**, Pair-matching performance landscape across binding event fluxes and unbinding rates for synthetic event pairs using the localisation precision of 100 kDa events. Accuracy is represented as opaqueness, decreases in accuracy attributed to falsely assigned pairs (false positives) and missed pairs (false negatives) are shown in red and blue respectively. Contours are drawn at accuracy values of 0.75, 0.6, and 0.5. Inset: to-scale representation of the outline of a β-galactosidase monomer (mass ∼116 kDa) and the localisation precision of a 100 kDa event. **d**, Localisation precision (scaled *α* =3.0) of all precursor (binding, blue) and product (unbinding/dissociation, orange) events from a β-galactosidase measurement plotted at the location in the FoV where they were detected; no pairing is applied. Inset: example event types with schematic illustration of behaviour. **e**, Combined mass histogram of 4 β-galactosidase measurements: precursor (binding) and product (unbinding/dissociation) events are shown as positive and ‘negative’ mass respectively, events successfully matched into pairs are shown in teal, unmatched events are shown in grey. Percentage matched represents the fraction of each class successfully matched. **f**, 2D spectrum of the absolute mass of matched precursor-product pairs from the same measurements. Each point represents one successfully matched precursor-product pair. Pairs on the diagonal correspond to unbinding and pairs below it to dissociation; pairs with m_product_ > m_precursor_ (red; 2.24%) are attributed predominantly to false-positive matches and lie above the false positive threshold (red dashed line). Marginal histograms are shown for precursor (binding, blue) and product (unbinding/dissociation) axes.

The major obstacle to reliable pairing of events is that drift of the glass coverslip over the course of a measurement (**Supplementary Fig. 1**) makes distinguishing a true paired outcome from a coincidental event in spatial proximity non-trivial. We developed a robust algorithm to extract and correct for the drift vectors of the glass, then quantified the resulting mass-dependent localisation precision. To do this, we simulated events across a range of masses by adding an experimental point-spread function (PSF) to experimental buffer backgrounds, and estimated the uncertainty in position by fitting a mathematical PSF model to the simulated signal (**Fig. 1b**). Examining the standard deviation of the resulting position-error distribution (*σ_p_* ) as a function of mass revealed (in line with interferometric contrast scaling with mass^8^) that higher-mass events have a higher signal-to-noise ratio, and *σ_p_* falls with increasing mass, following a reciprocal relationship (dashed line) up to ∼500 kDa, consistent with shot-noise-limited localisation scaling^22^. Above 500 kDa, *σ_p_* progressively deviates from this reciprocal scaling and plateaus at ∼0.9 nm; we attribute this floor to a residual non-shot-noise background^23^. Remarkably, above ∼240 kDa, the position error falls below the biomolecule’s minimum theoretical radius (gold line): for instance, the localisation precision of a 480 kDa complex lies comfortably within the outline of an example biomolecule of that mass (β-galactosidase, **Fig. 1b)**. This ability to super-resolve a biomolecule over extended time periods is the cornerstone of successful pair-matching.

We note that equivalent precision estimates for movements cannot be similarly obtained because their composite signature depends jointly on mass and displacement, which are effectively degenerate over regions of parameter space; we introduce a separate method to measure them below. Irrespective, the mass-dependent precision relationship sets the spatial scale for pairing events over time and defines how close a candidate outcome must lie to likely arise from the same precursor biomolecule. Guided by this relationship, we pair events by treating each localisation as a positional probability distribution of width *σ_p_* set by its mass, and assigning each product outcome to the candidate precursor binding event that maximised the overlap coefficient⁷ between the two distributions. As such we have developed means for multi-dimensional mass measurement of single molecules, as they undergo dynamic transitions, such as dissociation, on the glass surface. By analogy to eponymous mass spectrometry (MS) workflow, we term this approach tandem MP.

### Benchmarking the theoretical accuracy of tandem MP

To benchmark our approach, we simulated pairs of binding and outcome events at known positions and times, passed them through our tandem MP pipeline, and quantified the fraction correctly recovered. We mapped the accuracy (defined in **Methods**), for 100 kDa events (with localisation precision of 10 nm) as a function of binding flux (a proxy for analyte concentration) and unbinding rate constants (*k*_off_)(**Fig. 1c**). We chose to explore these parameters as they will affect the surface density of bound molecules and hence are critical for our ability to match events together correctly. For this simulation, we assumed perfect detection and fitting for events not in spatiotemporal proximity to another. We found that accuracy is primarily controlled by binding flux: at typical MP operating fluxes (≲0.5 events μm^-2^ s^-1^), accuracy ≳0.75 throughout the range 10^-3^ s^-1^ < *k*_off_ < 1 s^-1^. Two distinct failure modes bound the usable region: above *k*_off_ ≈ 1 s^-1^, binding and unbinding occur within the temporal resolution required to distinguish them as separate events, resulting in many missed pairs (false negatives) (**Fig. 1c, blue**). At fluxes >1 events μm^-2^ s^-1^, multiple candidate events fall within overlapping positional uncertainties and leads to an increase in incorrect matches (false positives) (**Fig. 1c, red**). Together, these define an operating window for reliable pairing: unbinding rates up to the temporal resolution of the instrument (∼3 s^-1^), at standard MP fluxes (≲0.5 events μm^-2^ s^-1^). Because *σ_p_* decreases with mass, this 100 kDa benchmark represents a conservative case, with events corresponding to heavier molecules tolerating correspondingly higher fluxes. For instance, accuracy increases by 5% from 100 kDa to 200 kDa at a typical MP flux of 0.5 events μm^-2^ s^-1^ and *k*_off_ of 0.3 s^-1^. We can also use our benchmarking to quantify the impact of our drift-correction and localisation-precision algorithms. Examining various binding-event fluxes and *k*_off_ values, we show both algorithms, when used together, significantly improve pair-matching accuracy (**Supplementary Fig. 2**). This two-thirds reduction in false positives is substantial because they represent the most pernicious source of error, and acts to realise the principle of faithful pair-matching of events in MP.

### Tandem MP reveals the dissociation pathways of a homotetramer

To test our concept for tandem MP, we first measured *E. coli* β-galactosidase, a homotetrameric enzyme with monomer mass of 116 kDa. After drift-correcting the frames of the video, we plotted each event recorded as a circle of radius 3*σ_p_* (i.e. ɑ=3.0, see **Methods**) based on our localisation precision relationship (**Fig. 1b**): with precursor binding events in blue, and product outcomes (unbinding or dissociation) in orange (**Fig. 1d**). 2165 events are distributed evenly across the field of view (FoV), with no evidence of preferential binding sites. Four patterns occur: 1) isolated blue circles (732 events, 33.8%), where molecules bound during the acquisition without subsequent detectable transitions; 2) isolated orange circles (140 events, 6.5%), where molecules bound before acquisition began, and thus only their departure was detected; 3) overlapping blue and orange circles (1019 events, 47.1%), where precursor binding events are followed by a co-localised unbinding or dissociation event; and 4) two overlapping blue discs (274 events, 12.7%), representing binding co-localised with another binding event (association). First, we tested whether these co-localised relationships are genuine. Randomising event positions in the β-galactosidase measurement (**Fig. 1d**) while retaining event mass and time provided a null rate for pair-matching (**Supplementary Fig. 3**). Precursor-to-product (binding-to-unbinding/dissociation) pairs were enriched 4.9-fold over null, compared with 1.1-fold for precursor-to-precursor (binding-to-binding) controls. The large enrichment of precursor-product pairs over null confirms that, at typical MP concentrations, the matched pairs reflect genuine precursor-product relationships rather than an artefact due to chance co-localisation.

We next combined pair-matching data from four measurements of β-galactosidase, and plotted the mass distribution of all detected events (**Fig. 1e, grey**), with precursor events as positive and product events as ‘negative’ mass. Precursor events are dominated by the intact tetramer at ∼480 kDa, with a minor free-monomer population at ∼120 kDa; product events resolve into monomer, dimer, trimer and tetramer species. Overlaying the events that pair successfully (**Fig. 1e, teal**) reveals precursor tetramers pair to product events at a high rate, whereas the monomer peak is almost entirely unmatched. This asymmetry indicates that tetramers bind to the surface, and either unbind intact, or lose one or more subunits, whereas a bound monomer (which has no subunit to lose) does not unbind.

Plotting the masses of each matched precursor-product pair separates three regimes by their position relative to the diagonal: intact unbinding (*m*_product_ = *m*_precursor_), dissociation (*m*_product_ < *m*_precursor_) and association (*m*_product_ > *m*_precursor_) (**Fig. 1f**). The upper-left region is sparsely populated (2.24% of pairs) and is consistent with the chance-assignment rate (**Supplementary Fig. 3**); we therefore treat it as residual false positives (using a linear threshold, defined in **Methods**) rather than genuine association. Pairs present along the diagonal represent a binding followed by a co-localised unbinding of equal mass, this region is dominated by tetramers binding and unbinding intact. The lower-right (dissociation) region is the most informative. Matched products form clusters at integer multiples of the monomer mass (∼120, ∼240, ∼360 kDa), and these originate overwhelmingly from intact tetramers, shown by the data falling almost entirely at a precursor mass of ∼480 kDa. Tandem MP therefore resolves the stoichiometry of subunit loss directly, distinguishing dissociation from unbinding and reporting how many subunits depart at each event. Our approach therefore reveals structure that is hidden in the standard one-dimensional mass histogram and emerges only in the paired precursor–product map.

***Tandem MP reveals differing disassembly pathways within a family of citrate synthases*** Having established that pair-matching resolves how complexes disassemble, we next applied it to a system with more complex oligomeric architecture to explore differences in behaviour. We examined citrate synthase (CS) from different organisms, *Se*CS, *Anc*CS, *Ds*CS and *Ni*CS.^17,18^ Respectively, these are a fractal-forming CS from *Synechococcus elongatus* (**Fig. 2a**) and an ancestral reconstruction (**Fig. 2b**) that are both primarily 18mers of ∼43 kDa monomers, and two hexameric examples, from *Desulfoluna spongiiphila* (monomer ∼50 kDa, **Fig. 2c**), and *Nitrospira inopinata* (monomer ∼42 kDa, **Fig. 2d**). Examining the mass spectra for each demonstrates that MP has sufficient mass resolution to distinguish the oligomeric states of all CS studied (**Supplementary Fig. 4**). *Se*CS displayed the richest dissociative behaviour (**Fig. 2a**): its 18-mer precursors (∼780 kDa) dissociated into a range of products: monomer, dimer, tetramer, hexamer and octamer, and also unbound intact. Smaller precursors showed their own pathways: hexamers dissociated into dimers or unbound intact, and dimers dissociated into monomers or unbound intact. *Anc*CS reproduced much of this behaviour but with clear differences (**Fig. 2b**): unlike the extant complex, few monomeric products were matched to 18-mer or hexamer precursors, and no 18-mer was observed to unbind intact, indicating a less stable assembly or stronger affinity to the glass surface. In contrast to *Se*CS, a significant portion of *Anc*CS hexamer precursors dissociated into tetramers. The non-fractal *Ds*CS bound predominantly as hexamers and dissociated mostly into dimers, with some tetrameric products; a small number of dimers and hexamers bound and unbound intact (**Fig. 2c**). In comparison, hexameric *Ni*CS (**Fig. 2d**) exhibited monomer and dimer dissociations but no tetrameric products; a small amount of monomer, dimer, and hexamer species unbound intact. This study reveals therefore that CS homologues that give very similar one-dimensional standard MP spectra (*Se*CS v *Anc*CS, and *Ds*CS v *Ni*CS) display distinct differences in the two-dimensional tandem MP plot.

**Fig. 2.**
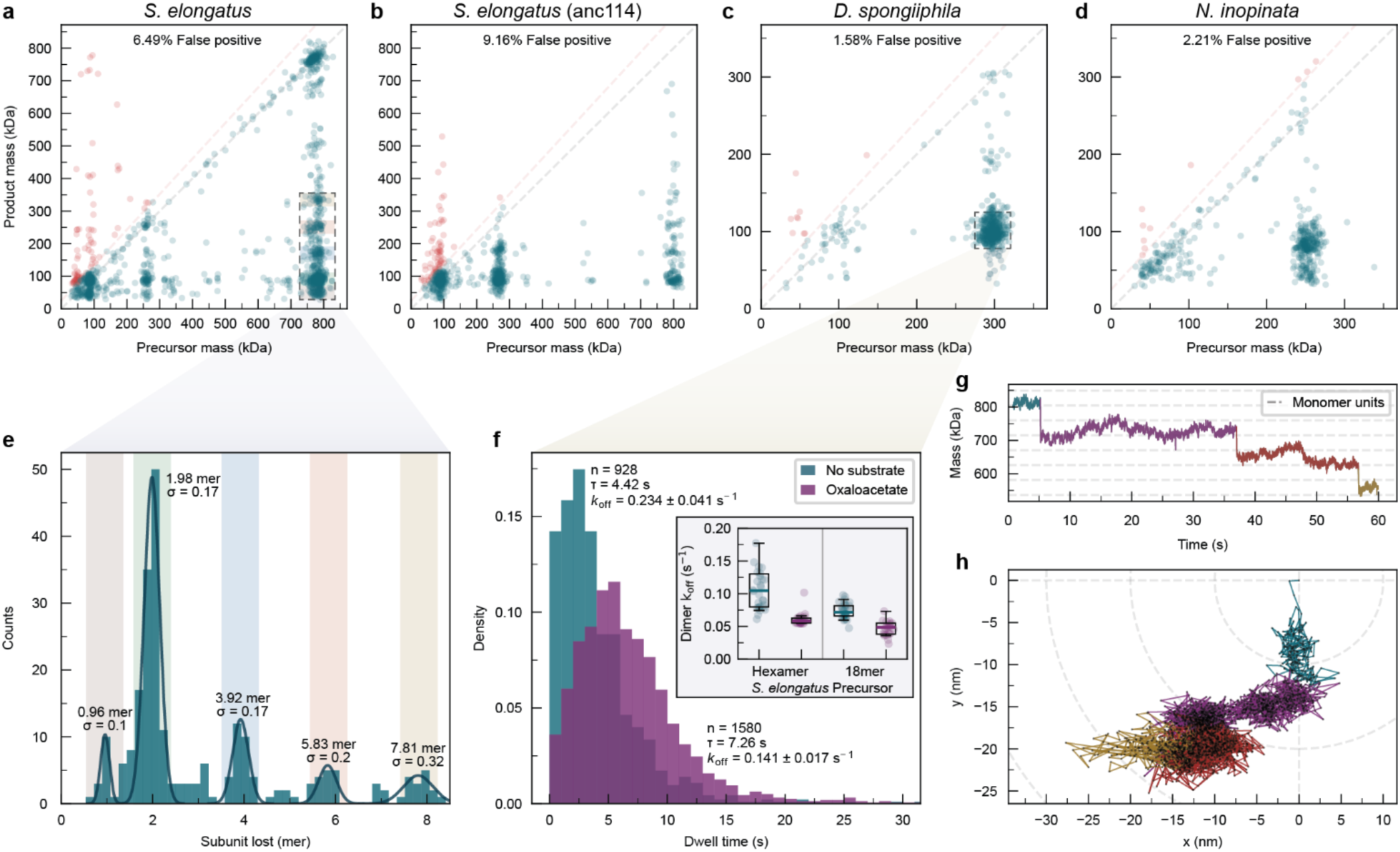
Pair matching resolves the oligomeric composition and disassembly kinetics of citrate synthases. **a-d**, Combined 2D spectrum of matched precursor-product pairs of : **a**, *S. elongatus* CS; **b**, its ancestral reconstruction (anc114); **c**, *D. spongiiphila* CS (no substrate); and **d**, *N. inopinata* CS. Lines and colours as in Fig. 1f; dashed boxes mark populations selected for analysis in **e** and **f**. **e**, Dissociation products paired to 18mer *S. elongatus* CS precursors with Gaussian fits to observed peaks at 1, 2, 4, 6 and 8 subunits, with 3, 5 and 7 largely absent. Oligomeric units calculated by dividing product mass *Se*CS monomer mass (44.7 kDa). **f**, Times between binding of *D. spongiiphila* CS hexameric precursors and first dimer dissociation, with substrate incubation (oxaloacetate, purple) and without substrate (teal), acquired under identical illumination conditions. Annotations show the number of dwell times measured (n), mean dwell time (*τ*), and effective dissociation rate constant (*k*_off_, with 95% confidence interval). Inset: equivalent dimer dissociation rates for *S. elongatus* CS hexamer and 18mer precursors. **g**, Mass over time trace of a dissociating *S. elongatus* CS 18mer complex obtained by fixed background continuous tracking (Fig. 3a). Segments are coloured by the dissociation detected. **h**, position over time trace of the same complex relative to first localisation.

### Tracking dissociation histories reveals structural organisation and dynamics of citrate synthases

Since tandem MP operates fundamentally at the single molecule level, it can also isolate the behaviour of individual complexes by linking multiple events to a single precursor, allowing us to follow the stepwise disassembly of individual molecules. Restricting the analysis to *Se*CS events that bound as intact 18mers, we first tallied every dissociation assigned to an 18-mer precursor (**Fig. 2e**). The mass lost per event is sharply quantised: peaks fall at 1.0, 2.0, 3.9, 7.8 subunits lost, corresponding to loss of a monomer, dimer, tetramer, hexamer, and octamer respectively. Therefore, the loss of subcomplexes with an even number of subunits is preferred (with some monomer dissociations also detected), revealing the dimeric substructure of *Se*CS. Next, linking successive events paired to the same individual precursor enables us to follow a single complex through its dissociation process. As an example, we can reconstruct the mass trace of a single 18-mer (a reconstruction rounded to the nearest oligomeric unit is show in purple), which sheds three successive dimers and then a hexamer over ∼10 s, resolving the disassembly pathway of an individual molecule step by step (**Supplementary Fig. 5)**. Following individual trajectories in this way presents tandem MP not just as an aggregate description but as a means for following the histories of individual biomolecules.

As each event carries a timestamp, the kinetics of disassembly is directly accessible. To explore the value of such measurements, we examined *Ds*CS, whose disassembly is dominated by a single dissociation (dimer products from hexamer precursors). We recorded the dwell time between a *Ds*CS hexamer landing and its first dimer dissociation, both in the presence and absence of its substrate oxaloacetate (**Fig. 2f**). The dwell time distributions for each condition can be summarised by a mean dwell time (*τ*) and an effective dissociation rate constant (*k*_off_ = 1/ *τ*). *k*_off_ decreases from 0.234 ± 0.041 s^-1^ without substrate to 0.141 ± 0.017 s^-1^ with substrate. These rates are within the operating window for reliable pairing (**Fig. 1c**), and allow us to quantify how substrate binding stabilises the hexamer against dimer loss. This demonstrates that dissociation constants can be obtained directly from tandem MP data, and prompted us to see if we could extract stoichiometry-specific dissociation constants. We therefore performed equivalent analysis of *Se*CS, quantifying *k*_off_ for both the hexamer and 18mer (**Fig. 2f, inset**). We find that dimers dissociate faster from the hexamer (0.105 ± 0.103 s^-1^) than the 18mer (0.072 ± 0.032 s^-1^). This is a surprising result given that there are more dimers available for dissociation from the latter. Furthermore, we found that the stabilisation conferred by oxaloacetate differed between the two stoichiometries. These experiments show how tandem MP can reveal how the kinetic stability of proteins is regulated by their assembly and interactions with small molecules.

### Laser-induced activation in tandem MP

Our inferral from tandem MP that *Se*CS is comprised of dimeric units is in line with the high-resolution structure and dimer-mediated assembly we previously reported^17^. Similarly, the reduction in its dissociation rate (and that of *Ds*CS) in the presence of oxaloacetate is in line with previous measurements on substrate-induced stabilisation of CS^24^. This suggests that tandem MP data, despite reporting on the behaviour of molecules bound to the glass surface of the cover slip, encodes structural and dynamical signatures of the native protein complexes. This important observation invited us to investigate the mechanism by which dissociation occurs. We first noted that the dwell time distributions are peaked rather than the single-exponential shape expected from a simple first-order dissociation step. This is consistent either with a multi-step process^25^, or time-dependent reaction kinetics, such as progressive heating increasing the rate of thermally activated dissociation during the observation period^26^. The latter is intuitively plausible in our experiment and analysis. In MP, the laser is focussed to the glass-water interface, and therefore in conventional MP analysis, a protein is exposed only momentarily (∼100 ms) to the full intensity of the laser (10^2^ -10^3^ kW cm^-2^)^8^, as it binds to the surface and is measured (once only, by ratiometric imaging). However, while adsorbed to the surface, the protein remains continuously exposed to the MP laser, for seconds or longer. To test this hypothesis, we examined whether the dissociation behaviour we see is dependent on illumination wavelengths and powers (**Supplementary Table 1**). We find that *k*_off_ is highly laser dependent, with shorter wavelengths and higher powers leading to faster dissociation. This suggests that the mechanism of tandem MP is laser-stimulated dissociation due to non-resonant heating at the glass-water interface.

### Tracking mass and position changes validates tandem MP results

Next, we wished to validate that we were genuinely observing single particles undergoing multiple dissociation events. We therefore asked whether continuous particle tracking^27^ could recover the full trajectory between paired events. In ratiometric imaging, only temporal changes are visible, revealing a molecule only as it binds, unbinds, moves, or dissociates; once bound and stationary, it disappears from the image (**Fig. 3a, top row**). Creating a ratiometric view with the background fixed temporally before a particle binds retains its signal continuously; with drift correction applied to the native frames, this can yield its mass and position over time. Applied to *Se*CS, the mass trace of an 18-mer shows sharp, step-like decreases quantised in monomer units (**Fig. 2g**), matching the dissociative behaviour characterised above by means of tandem MP. Furthermore, on rare occasions, we observed dissociation products rebinding near the precursor *Se*CS complex (**Supplementary Fig. 6**), further supporting the precursor-product relationship. The same particle also translates across the surface over the 60 second trace (**Fig. 2h**). Importantly, the fact that mass and position changes are not strictly correlated means that the mass changes observed are not an artefact due to a change in the underlying glass background. In addition, this nanoscale movement exceeds the ∼15 nm diameter of the complex^17^, so cannot be attributed to a shift in centre of mass due to subunit loss. Our mass tracking data therefore reveals directly how individual molecules undergo dissociation events on the surface, and validates our tandem MP insights.

**Fig. 3.**
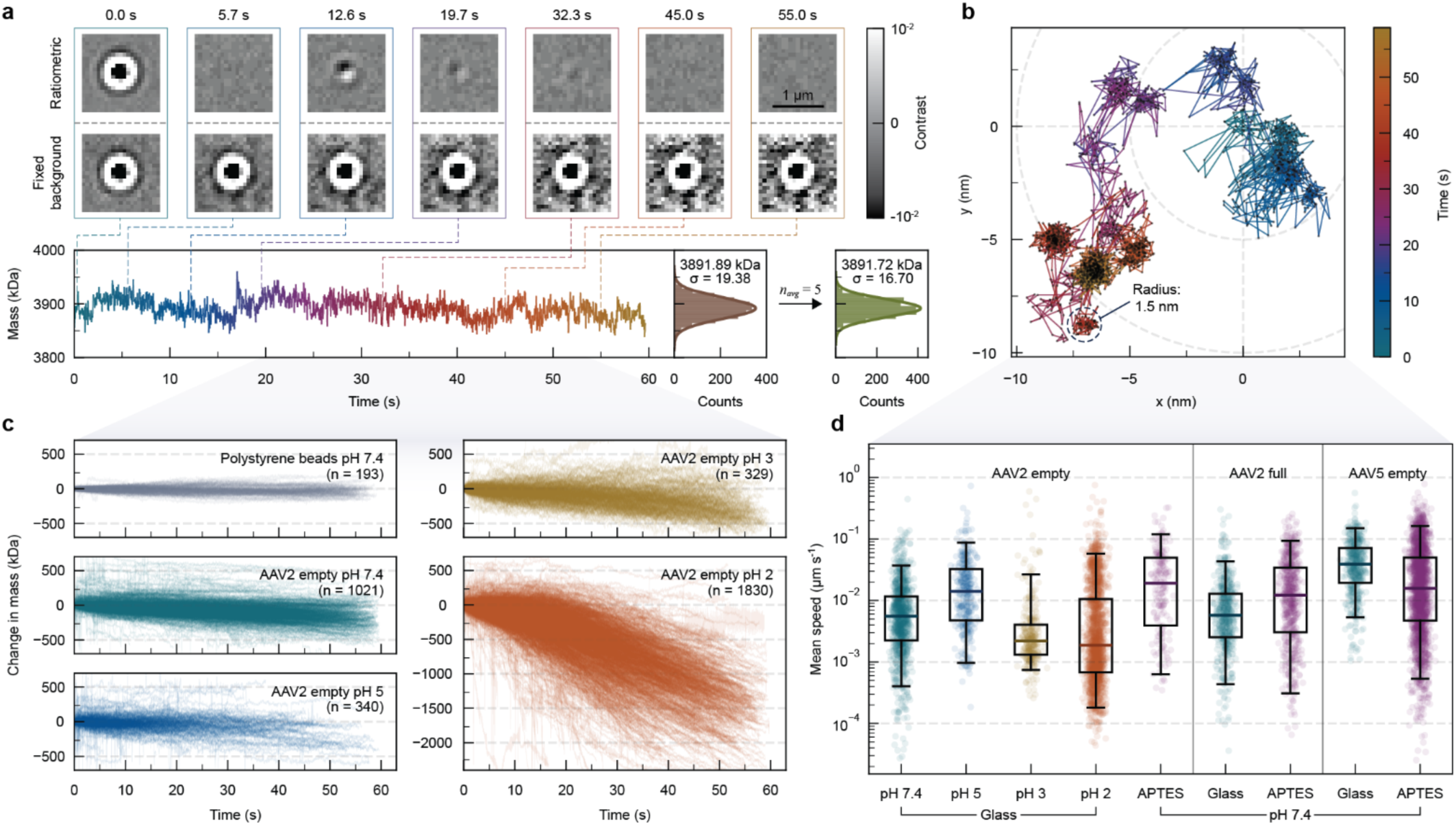
Fixed background particle tracking reveals the dynamics of AAV capsids at the single-molecule level. **a**, Mass over time trace (bottom) of an empty AAV2 capsid (glass, pH 7.4) with corresponding ratiometric (top) and fixed background (middle) images. The capsid is visible transiently in ratiometric images upon binding (0.0 s) and movement (12.6, 19.7, 32.3 s), but remains visible throughout in the fixed-background images. Right, mass distributions with Gaussian fit before and after averaging over *n_avg_* = 5 frames. **b**, Position over time trace of another empty AAV2 capsid (glass, pH 7.4) relative to its first localisation, coloured by time. Dashed circle, significant-movement threshold (radius 1.5 nm) from simulations (**Supplementary** Fig. 10). **c**, Change in mass relative to the first frame for polystyrene beads (20 nm) and empty AAV2 capsids on glass at pH indicated. Each line represents the change in mass of one tracked capsid. **d**, Mean movement speed of tracked AAV capsids under various conditions, calculated using a significant movements threshold (1.5 nm, shown in **c**). Empty AAV2 capsids on glass at pH 7.4 (teal), 5 (blue), 3 (yellow), and 2 (orange); and on APTES functionalised glass at pH 7.4 (purple). Full AAV2 capsids at pH 7.4 on glass (teal) and APTES (purple). Empty AAV5 capsids at pH 7.4 on glass (teal) and APTES (purple). Each point represents the mean movement speed of one tracked capsid.

### Continuous tracking of single AAV capsids improves precision and reduces error of mass measurement

Having established the feasibility of measuring the particles on the glass surface for extended periods of time, we reasoned that this capability should allow improved mass measurement. To test this, we turned to AAVs, viral capsids that are visibly mobile on glass in MP measurements, and whose large mass (>3MDa) gives excellent signal-to-noise ratios, even with the accumulating background that arises from a fixed background (**Fig. 3a**). Monitoring a representative AAV2 capsid over 60 s, from the moment of binding (0.0 s) reveals several distinct movement events (at 12.6, 19.7 and 32.3 s) as transient signals in the ratiometric image. Instead, in differential imaging with a fixed background, the capsid remains visible throughout, yielding a continuous mass trace around 3900 kDa, allowing its mass to be determined with high precision (**Fig. 3a**).

Continuous tracking can also return the capsid’s trajectory on the surface with nanometre-level precision (**Fig. 3b**), in-line with expectations from our benchmarking (**Fig. 1b**). The trajectories show that the AAV explores a small number of discrete sites rather than diffusing freely, which may reflect the capsid rocking between distinct contact points on its surface or glass inhomogeneities at the nanoscale (**Fig. 3b**). One benefit of particle tracking is that movements are accommodated for: in standard MP analysis, unless they filtered out effectively, movements give rise to an artefactual pair of symmetric peaks about zero contrast (**Supplementary Fig. 7, grey**). These arise because a movement event is erroneously fitted as adjacent binding and unbinding events of equal and opposite contrast, the bright and dark lobes of a single movement (**Fig. 1a**), resulting in PSF fits with elevated errors. Particle tracking instead resolves each movement into its components (co-located unbinding and binding events) driving fit errors an order of magnitude lower (**Supplementary Fig. 7, teal**) and improving per-particle mass precision by measuring the same scatterer multiple times.

### Resolving the dynamics of AAV capsids at the single-particle level

We next aimed to explore to what extent the mass and position changes of AAV might reflect the surface properties and stability of the particles. Several AAV serotypes exist, and they have differing surface properties, and undergo pH-dependent dissociation. We therefore chose to monitor thousands of AAV capsids across a range of physiologically relevant pHs and different surfaces to follow their behaviour in real time. As a baseline, we first tracked polystyrene beads, which have a similar refractive index^28^ to protein complexes but are more thermostable. As expected, their mass traces stayed flat over the 60 s measurement (**Fig. 3c**, **Supplementary Fig. 8, grey**). By contrast, empty AAV2 capsids at neutral pH showed a small but steady mass decrease (**Fig. 3c, teal**), consistent with dissociation of the capsid and quantifiable as a net-negative decay rate from a linear fit to each trace (**Supplementary Fig. 8, teal**). This decay was essentially unchanged at pH 5 and pH 3 (**Fig. 3c, Supplementary Fig. 8, blue and yellow**). The insensitivity across three pH units is consistent with dissociation being laser-induced, consistent with our tandem MP mechanism investigations above (**Supplementary Table 1**), instead of a pH-induced effect due to being in the immediate vicinity of the negatively charged glass surface. However, at pH 2, a sufficiently acidic condition to stimulate genome release from AAVs in the lysosome^29^, the mass decrease became substantially steeper, consistent with acid-induced capsid dissociation (**Fig. 3c & Supplementary Fig. 8, orange**). Examination of the mass decay curve reveals clear peaks corresponding to intermediate states, in this case separated by ∼60 kDa steps, attributable to the loss of viral coat protein (VP) monomers (**Supplementary Fig. 9**).

The same experiments also report each capsid’s trajectory on the surface, from which we could quantify a mean speed. We considered only movements >1.5 nm as significant (**Fig. 3b**), based on our simulations (**Supplementary Fig. 10**). By quantifying the AAV trajectories across the different conditions, we found that the mean speed varied significantly (**Fig. 3d**). Lowering pH from 7.4 to 5 increased the mean speed, coinciding with the pH range at which VP1u externalisation is reported^30,31^ (**Fig. 3d, teal and blue**). At the lowest pHs, the mean speed was at its slowest, attributable to increased capsid–glass affinity as the capsid becomes more protonated (**Fig. 3d, yellow and orange**). Comparing surfaces at neutral pH, both empty and full AAV2 capsids moved faster on APTES (positively charged, **Fig. 3d purple**) than on glass (negatively charged, **Fig. 3d teal**); but we could not distinguish empty from full AAV2 capsids by movement under our conditions. Interestingly, we noted differences between the two serotypes were identifiable: empty AAV5 capsids moved faster than AAV2 on glass and, unlike AAV2, slowed on APTES. This may reflect the different surfaces properties of the two capsids^32^, with AAV5 lacking the cationic heparan-sulphate binding patch on AAV2^33^. Differences between surfaces and serotypes were not visible in the capsid mass traces (**Supplementary Fig. 11**), demonstrating the orthogonality of tracking position. Together these experiments demonstrate that proteins can be mobile on the glass surface, with their motion reflecting their surface properties, and how they interact with the charges on the glass surface.

## Discussion

Here we have presented two analysis modalities for MP beyond the counting of independent landing events: tandem MP and particle tracking. The approach relies our ability to localise the position of particles to a precision well below the Abbé diffraction limit of the MP laser light (174 nm on a TwoMP, Refeyn Ltd). This is made possible by correcting for the lateral drift of the coverslip glass during a measurement, and allows us to link, with tolerable ambiguity, each binding event to its subsequent fate at the same location on the surface. Our Python code is freely available, and useable on MP data collected on commercial instruments with minimal alterations to experimental protocols. The software enables the transformation of the typical ratiometric MP histogram, in which each particle is measured only transiently, with continuous, resolved single-particle histories.

The tandem MP modality bears parallel with tandem MS of protein complexes (in the form of ions in vacuum) where precursors are dissociated to reveal their subunit composition^34^. The tandem MS workflow is performed by selecting protein complexes of a certain mass (technically, mass-to-charge ratio), and then intentionally activating them, typically either by collisions with gas, electrons, or laser photons, before measuring the products. In tandem MP, no such selection is required as we are dealing with molecules one at a time. Dissociation in tandem MP is unintentional: it is a combination of the natural dissociation kinetics of the protein, augmented by heating at the glass-water interface, a consequence of continuous irradiation by the MP imaging laser. An advantage of tandem MP over tandem MS is that the dissociation occurs in solution, albeit at the glass surface. The dissociation pathways of desolvated protein complexes in tandem MS frequently do not represent the native structural organisation^35,36^, whereas our experiments on CS and AAVs demonstrate that the dissociation pathways operative in MP reflect the pathways in solution^17,18^. Aside from revealing the units of assembly, our tandem MP measurements of CS also uncovered disassembly pathways that are invisible in the standard one-dimensional mass spectrum, as well as the associated kinetics.

Particle tracking is a logical extension of the pair-matching principle that underpins tandem MP, and follows the mass and position of a single persistent particle continuously over its lifetime on the surface. We were able track particles on glass and account for the movement-induced artefacts that distort standard analysis. Examining the trajectories of AAV capsids, at different pHs and upon varying the charge of the glass surface, showed that mass and position readouts are orthogonal and can be used to report on surface chemistry as well as dissociation behaviour.

Our approach converts a standard MP measurement from a static, population-level measurement into a time-resolved, massively parallel, single-molecule interrogation. The ability to do so with minimal adjustment to the MP workflow, and not requiring the addition of labels or modification of the glass surface makes the measurement immediately accessible. However, the approaches also have limitations: both tandem MP and particle tracking are concentration limited, with the former having a greater concentration tolerance than the latter; pair-matching accuracy falls with event flux, while particle tracking relies on a fixed background image that requires molecule’s scattering signal to be spatially resolved. Similarly, both methods are mass limited: pair-matching is theoretically possible at the MP detection limit however low localisation precision and low detection efficiency of MP at small masses can introduce false positives; results in this regime should be treated with care. In particle tracking, despite dynamic background frame selection, the accumulation of noise over time limits the sensitivity for low mass molecules, and small changes in a species’ mass or position. Lastly, both methods are censored by the integration time used, which sets a floor on how quickly a mass or position change event can be detected, and the acquisition endpoint; both can be tuned depending on the system being studied.

Our approach is highly complementary to the MP approach in which molecules confined or attached to lipid bilayers are monitored^13–15^. In the bilayer setup, the dynamics of membrane associated proteins can be interrogated in a relevant environment^13–15;^ alternatively a soluble protein can be immobilised to the bilayer via a tag and suitably modified lipid^37^. While experimentally more involved, this strategy leads to the proteins being held a short distance (defined by the linker between protein and tag) from the surface, giving the protein more freedom of movement that if bound to directly to glass. This reduced steric hindrance is likely to return more representative measurements of the absolute timescale of molecular transitions, in particular association reactions^16^. In addition, the free diffusion of the attached protein in the lipid bilayer means that it samples much of the imaging surface, acting to average out local differences in reflectivity that lead to apparent heterogeneities in mass measurement^23^. Conversely, the advantage of the approach presented in the present work lies primarily in its simplicity, but also in the much larger number of particles that can be monitored in parallel, readily allowing ∼100 molecules to be tracked in a single 60s movie. This could be particularly well suited to single-molecule immunoassays, where the surface is patterned with many binding sites in parallel^38^.

Throughout our investigations, we note that dissociation behaviour varies with illumination conditions (**Supplementary Table 1**) and displays kinetic features consistent with non-isothermal heating (**Fig. 2f**). We therefore consider the dissociation of molecules on the surface to most likely be accelerated by local heating of the interface by the MP laser. We had considered that the acidic pH very close to the glass-solution interface might influence the dissociation behaviour of the bound proteins. However, given the Debye length in typical buffers, this low pH regime extends < 2nm from the surface, which is small on the scale of the proteins detectable by MP. Indeed, performing experiments at increasingly acidic pHs (**Fig. 3c**) did not display a monotonic increase in dissociation rate that would be expected were surface pH effects to be operative. Nevertheless, we believe that surface modification is a fertile area for future control of dissociation behaviour, particularly as not every protein complex will display rich dissociation behaviour. With this and other opportunities to engineer further control of activation, we believe that the work presented here represents only a small subset of the opportunities afforded by tracking particle histories in MP.

## Methods

### Experimental

All data was acquired on a TwoMP (Refeyn Ltd.), unless stated otherwise. For pair-matching, regular field of view (10.9 × 4.3 μm^2^, 900 × 354 pixels) was used with a frame rate of ∼500 Hz, frame binning of 10, and pixel binning of 6 × 6. For particle tracking, large field of view (16.9 × 12.1 μm^2^, 1398 × 996 pixels) was used with a frame rate of 105-132 Hz, frame binning of 3, and pixel binning of 6 × 6. A post-binned pixel size of 72.6 nm was used for unit conversions. The acquisition image, acousto-optic deflector run times, focus position stabiliser, and autofocus image were calibrated at the start of every measurement session. The instrument was calibrated at least once during every measurement session using a commercial (MassFerence^TM^ P1, Refeyn Ltd.) or in-house protein standard.

Glass coverslips (Menzel-Gläser, 1.5 H, 24 × 50 mm) were cleaned by sequential sonication in 50% isopropanol/MilliQ water and MilliQ water (10 mins each) and dried under a nitrogen stream. For APTES functionalisation, a modified version of the protocol developed by Kratochvíl et al.^23^ was used. Briefly, plasma-cleaned coverslips were functionalised with APTES by dipping in 2% (v/v) APTES in HPLC-grade acetone for 5 min, with 1 min of sonication in this solution. Coverslips were then transferred to fresh HPLC-grade acetone and sonicated for 1 min, followed by transfer to MilliQ water with several washes and a further 1 min sonication. Functionalised coverslips were dried under a stream of nitrogen and baked at 110 °C for 1 h before use.

Reusable silicone gaskets (Grace Bio-Labs CultureWell^TM^, 3 mm diameter, 1 mm depth) were placed on clean coverslips and placed onto the stage of the mass photometer with immersion oil (Zeiss Immersol^TM^ 518F) covering the objective. All measurements were acquired for 60s.

#### Protein

Protein samples were diluted to form a 1 μM stock using PBS and further diluted in-well using PBS or a 500 μM oxaloacetate-in-PBS solution (filtered through a 0.22 µm PVDF filter; Millex^®^-HV, Merck Millipore Ltd.). Final in-well concentrations were 2–6 nM for particle tracking measurements and 15–50 nM for pair-matching measurements.

#### AAVs

AAV capsid standards (PROGEN Biotechnik GmbH) were buffer-exchanged by the addition of 200 µL PBS followed by centrifugation for 5 min at 4000 rpm using 100 kDa MWCO centrifugal filters (Amicon^®^ Ultra-4, Merck Millipore Ltd.). Samples were then diluted 50-100× in-well using PBS adjusted to varying pH (using 6 M HCl) which was filtered (0.22 µm PVDF filter; Millex^®^-HV, Merck Millipore Ltd.) prior to use.

#### Polystyrene beads

Polystyrene bead size standards (22 ± 2 nm diameter; 3000 Series Nanosphere™, Fisher Scientific UK Ltd) were prepared according to the manufacturer’s protocol. Bead suspensions were gently inverted and sonicated for 10 s, immediately followed by dispensing of 2–3 drops into 10 mL PBS. The resulting suspension was diluted in PBS to a suitable concentration prior to measurement.

### Drift correction

The drift offset (*^δ_t_*) between a baseline frame (*B_t_*) and a target frame (*I_t_*) can be estimated by minimising the root-mean-square error (RMSE) between the target frame and a shifted baseline.

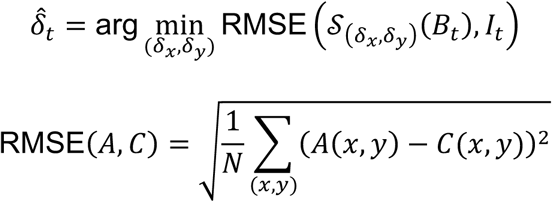

Where S_(ẟ_*_x_*_,ẟ_*_y_*_)_ denotes the translation of a frame by (*δ_x_*, *δ_y_*) pixels, implemented via cubic spline interpolation^39^, the RMSE is restricted to the region of overlap between target and baseline frames. The minimisation is performed by differential evolution^39^ over a bounded search domain (*δ_x_*, *δ_y_*) ∈ [−6, 6]^2^.

A drift corrected target frame (*I*^B^*_t_*) can be obtained via the below relation.

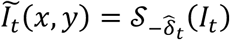

For an MP movie, the baseline frame and all corrected frames were subsequently cropped to a single common region-of-interest, defined as the area remaining valid across the full range of estimated drift for the entire movie. An example glass drift vector is shown in **Supplementary Fig. 1**.

### Pair-matching

The FoV coordinates (*x*, *y*), frame index (*i*), and mass (*m*) of ratiometric events arising from scatterers binding (*b*) and unbinding (*u*) from the glass-water interface were obtained from drift corrected MP movies using DiscoverMP (v2025 R1, Refeyn Ltd.), threshold 2 (under advanced mode analysis settings) of 1.5 was used to discard movement events from analysis. The mass dependence standard deviation of position error (*σ_p_*(*m*)) for ratiometric events was modelled by least squares minimisation to data in **Fig. 1b**, this was floored at 1 nm to prevent unrealistic small estimates for high-mass particles.

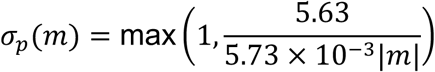

For each unbinding event, the binding event maximising the positional overlap probability^40^ (*P_ub_*) was selected as its match, subject to a minimum-overlap acceptance threshold:

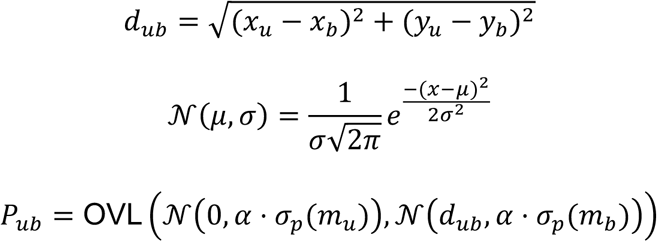

Where *d_ub_* represents the spatial distance between binding events and candidate unbinding events (*i_u_* > *i_b_*) and *α* denotes a scale factor applied to the mass-derived positional uncertainty. For all pair matching analysis, a scale factor of *α* = 3.0 was used and a minimum overlap threshold of *P_min_* = 0.05 was used to filter out unlikely pairs.

At typical MP concentrations we expect precursor-precursor pairs (or association) to be extremely rare (**Supplementary Fig. 3**), we therefore treat observed associations as false positives in the matching process. A pair is flagged as a false positive according to the below criteria, which accounts for the detection limit in MP and the increase in peak width with mass.

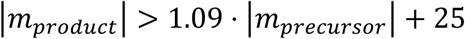

### Particle tracking

Particle tracking analysis was performed on the fixed background image using an in-house Python package, available on request. Briefly, for an identified ratiometric event arising from a scatterer binding to the glass-water interface (*b*) occurring at frame index (*i_b_*), a background thumbnail image (*B*(*x*, *y*)) for the area of the FoV in which the event takes place is calculated from the drift corrected MP movie (*I_t_*).

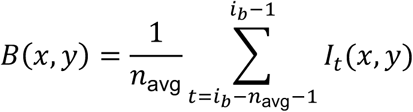

Where *n*_avg_ represents the number of frames averaged to form a background. The fixed background image (*F_t_*(*x*, *y*)) is obtained by dividing every subsequent drift corrected frame (*t* ≥ *i_b_*) by the background image.

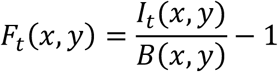

The fixed background image was computed for every frame where the scatterer is bound to the glass-water interface. Contrast, mass, and position values were estimated at every frame by minimising the sum of squared residuals between the signal arising in *F_t_* and a mathematical PSF. Tolerances and thresholds were used to account for and exclude other scatters in close spatiotemporal proximity to the scatterer of interest.

### Simulation

#### Mass-σ_p_ relationship

Experimental PSFs were obtained by cubic spline interpolation^39,41^ of pixel-wise median composite of high-SNR (> 95^th^ percentile) landing events. Experimental PSFs of known simulated mass were superimposed onto real, buffer-only background sub-stacks (randomly drawn experimental buffer movies) at a fixed central sub-pixel position (0,0). A mathematical PSF was fit to the resulting ratiometric image by minimising sum-of-squared residuals to recover the sub-pixel position (*δ_x_*, *δ_y_*), which is also the position error. This was repeated 1,000 times per mass with each repeat drawing from an independent background. Standard deviation (*σ*) of the position error distribution at every mass is computed by fitting a single spherical 2D Gaussian. The theoretical minimum radius for a globular protein of a given mass was calculated using a specific volume of 0.7446 mL/g ^8^.

#### Pair-matching performance benchmark

The position of synthetic ‘pair’ (binding then unbinding) events was uniformly simulated within FoV bounds, normally distributed position noise (N(0, *σ_p_*(*m*))) was added to each event independently according to the model outlined above. The temporal offset (Δ*t*) between the binding and unbinding events, or dwell time, of a pair was drawn from an exponential distribution parameterised by the unbinding rate (*k*_off_), Δ*t* ∼ *e^k^*^off^ . The number of simulated binding events were set from a target areal event rate: *n_events_* = *ρ* ⋅*A*_FoV_ ⋅ *T*, where *ρ* is the event density, *A*_FoV_ is the FoV area, and *T* is the total duration of the simulation. Both *ρ* and *k*_off_ were varied independently to construct a 2D performance landscape. Events occurring in close spatiotemporal proximity were merged. The remaining events were matched using the same overlap-probability pair-matching procedure described above (*P_ub_*, *P_min_* = 0.05, *α* = 3.0).

A match was scored as a true positive (*TP*) if the matched binding and unbinding events originated from the same simulated pair, and a false positive (*FP*) otherwise (mismatched pairing); false negatives (*FN*) were calculated as the number of ground-truth pairs not recovered as true positives. Ground-truth pairs whose true dwell time fell within 3 frames of the binding event (indistinguishable from simultaneous merging) were excluded from the ground-truth pair count. Accuracy was calculated as:

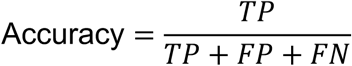

## Code availability

The Python software for tandem MP and particle tracking is available on GitHub: https://github.com/BeneschGroup/TandemMP

## Acknowledgements

The authors would like to thank all technical and research support staff who make our research possible.

## Author contributions

Conceptualisation: A.R.O, J.L.B, and J.L.P.B. Investigation: A.R.O, J.L.B., J.B., M.P., J.S.P., M.S., T.T. Resources: F.L.S. Formal analysis: A.R.O and J.L.B. Software: A.R.O., J.L.B., M.F.H. Writing – original draft: A.R.O, J.L.B., and J.L.P.B. Writing – review & editing: all authors. Funding acquisition and supervision: T.T., S.U., G.K.A.H., P.K., and J.L.P.B.

