## Supplementary information for "Super-resolved mass photometry reveals the dissociation pathways of individual protein complexes"

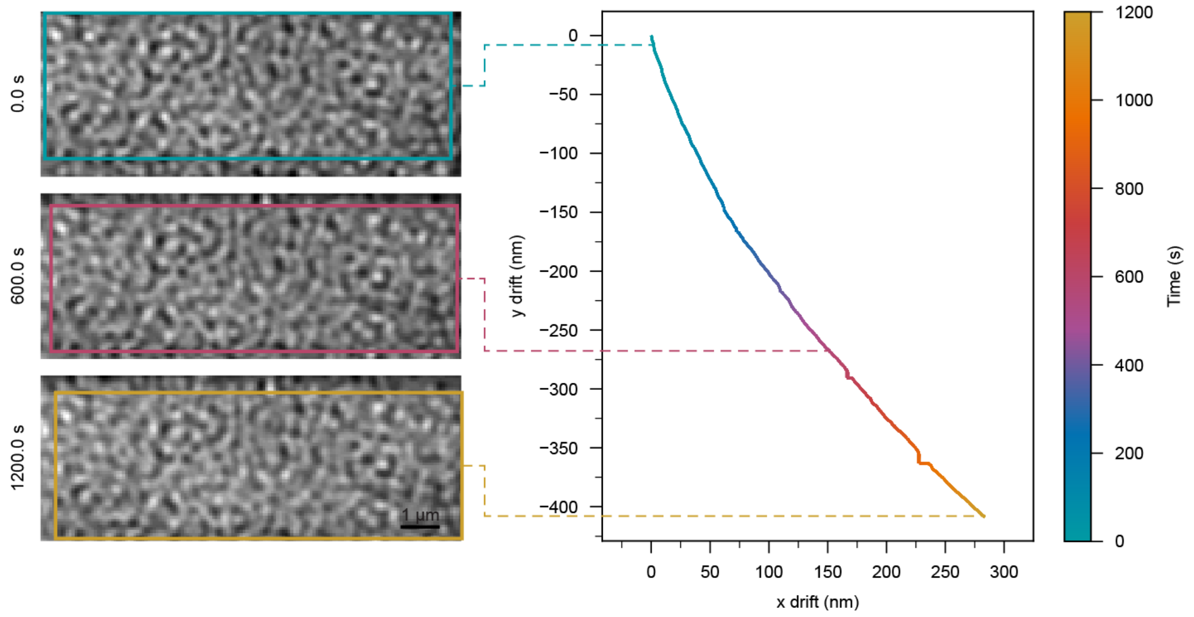


**Supplementary Fig. 1.** Lateral drift of the coverslip over a 20 minute measurement. Left, native (non-ratiometric) frames at 0, 600 and 1,200 s, showing the scattering pattern from glass roughness. Coloured boxes mark the region of glass that remains within the field of view throughout the measurement, positioned in each frame using the measured drift. Right, cumulative coverslip drift relative to the first frame, obtained by via drift correction (see **Methods**) and coloured by time. The coverslip drifts ~500 nm over 20 minutes or ~25 nm over a 60-s acquisition, well above the localisation precision of most protein complexes detectable in MP (**Fig. 1b**) and the capsid movement threshold (**Fig. 3b**).

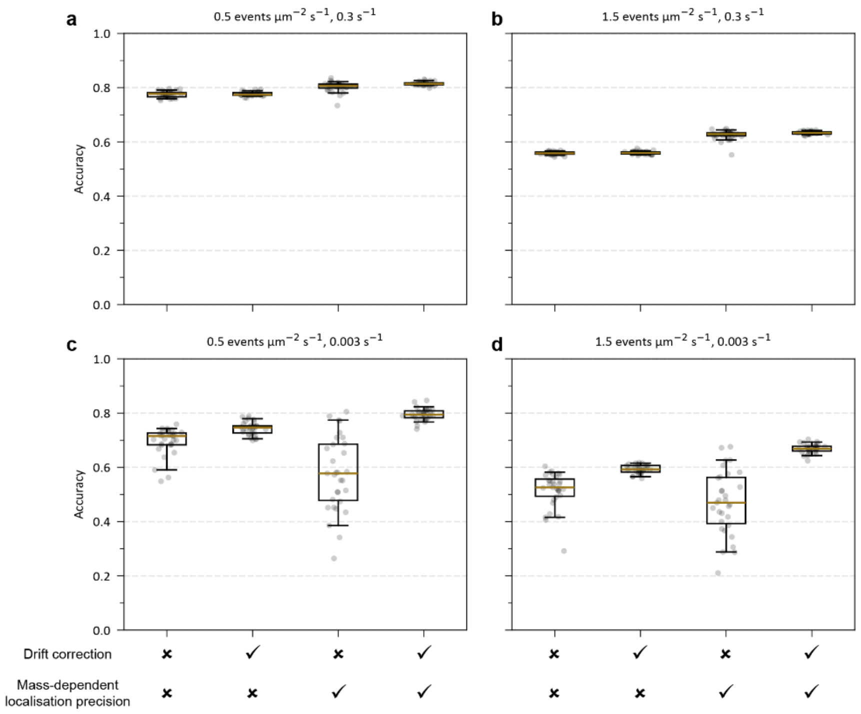


**Supplementary Fig. 2.** Drift correction and mass-dependent localisation precision improve pair-matching accuracy. Accuracy of pair-matching synthetically generated pairs at various binding event fluxes and unbinding rates ($k_{\mathrm{off}}$). Synthetic pairs were generated across a uniformly sampled range of masses from 50 to 1,000 kDa (unlike **Fig. 1c** which uses constant mass of 100 kDa) then analysed with or without drift correction and with either mass-dependent localisation precision or a fixed precision of 10 nm (equivalent to 100 kDa, **Fig. 1b**). Each point represents the average of 3 repeats of a simulation using a drift vector obtained from a real measurement.

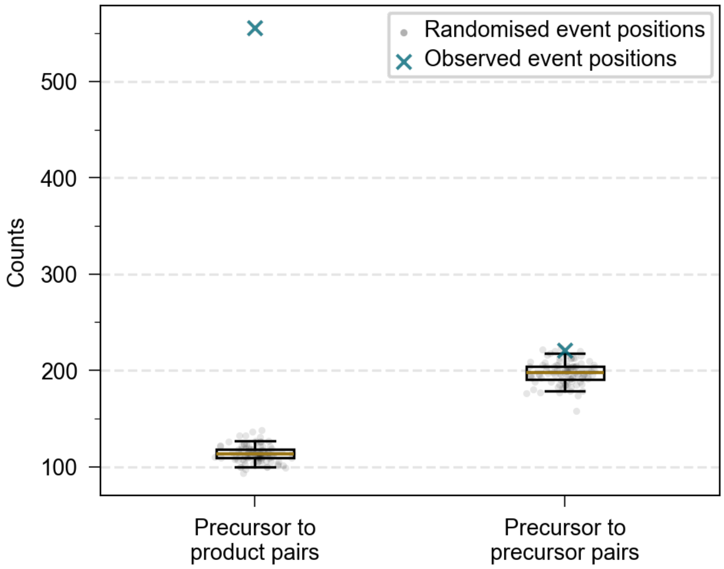


**Supplementary Fig. 3.** Matched precursor–product pairs are enriched over chance co-location. Number of successfully matched precursor-product (binding-unbinding/dissociation) and precursor-precursor (binding-binding) pairs from the β-galactosidase measurement shown in **Fig. 1d**, using event positions observed in the measurement (teal crosses) and randomised event positions (grey dots, 100 repeats). Pair matching was applied identically in both cases; only event positions were randomised, the mass and time of events was retained from the β-galactosidase measurement.

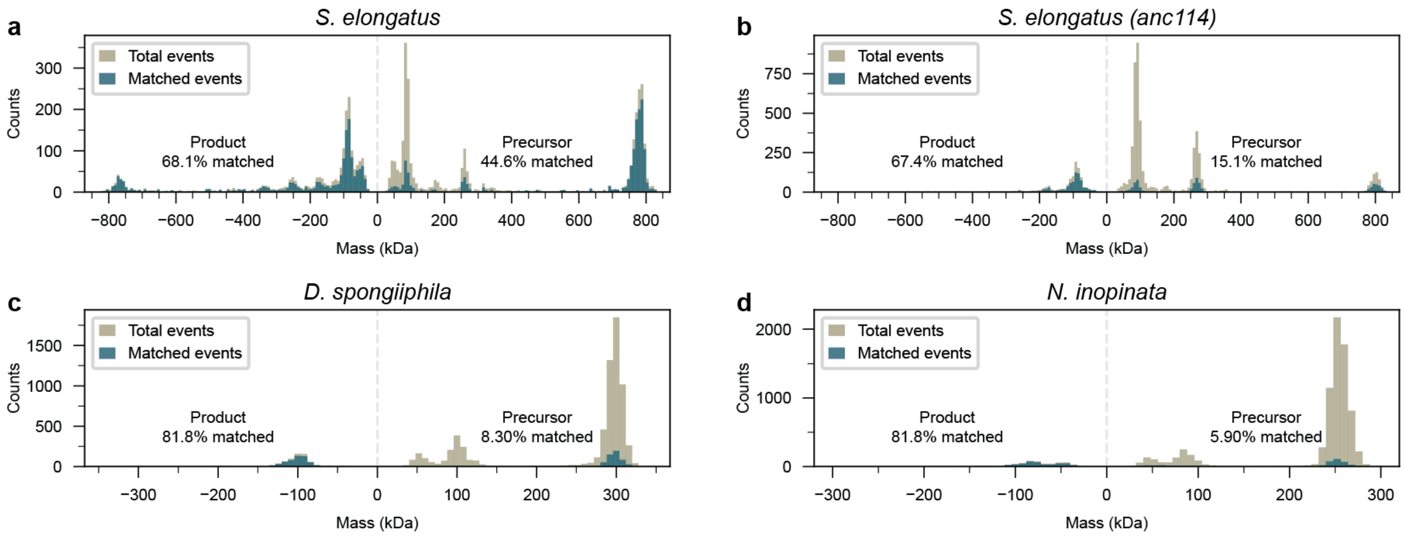


**Supplementary Fig. 4.** Landing assay mass distribution for the citrate synthase constructs: **a**, *S. elongatus* CS; **b**, its ancestral reconstruction (anc114); **c**, *D. spongiiphila* CS; **d**, and *N. inopinata* CS. Precursor (binding) and product (unbinding/dissociation) events are shown as positive and 'negative' mass respectively, events successfully matched into pairs are shown in teal, unmatched events are shown in grey. Percentage matched represents the fraction of each class successfully matched. Matched events correspond to the pairs shown in **Fig. 2a-d**.

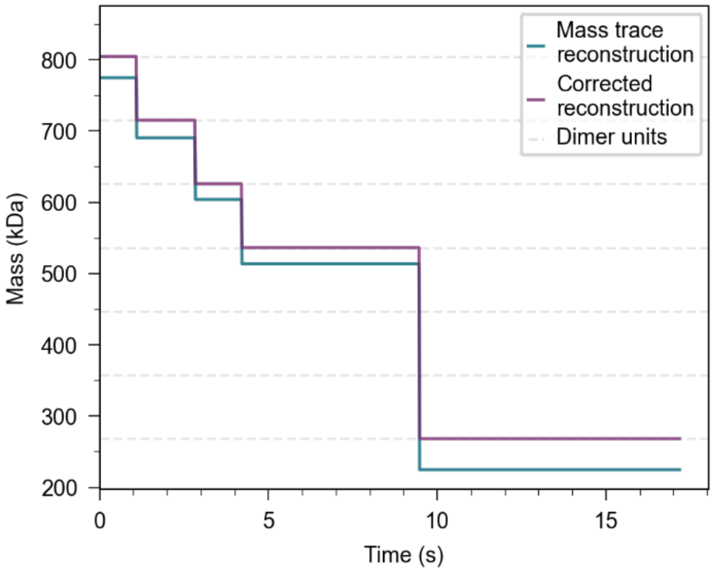


**Supplementary Fig. 5.** Pair matching reconstructs the fate of a single *S. elongatus* CS 18-mer. Mass over time of one 18-mer, reconstructed by linking all product events matched to a common precursor binding event in time order. Teal, uncorrected reconstruction; purple, reconstruction with plateau rounded to the nearest oligomeric state.

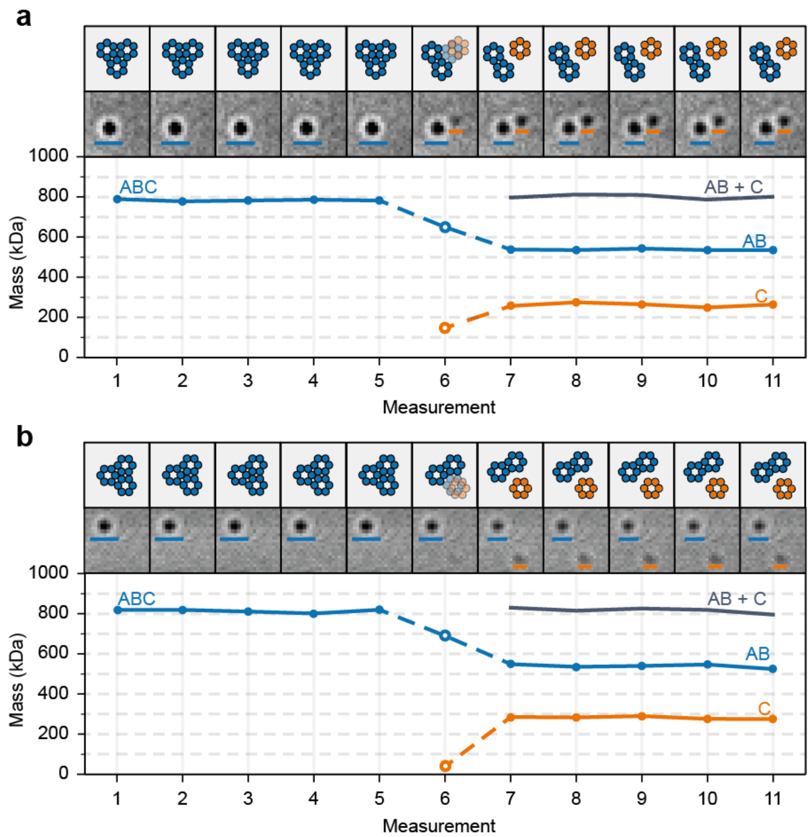


**Supplementary Fig. 6.** Two examples (**a** and **b**) of a hexameric product originating from a 18mer *Se*CS precursor adsorbing to the glass surface near the precursor complex. Top row, schematic of the precursor (blue) and product (orange) complexes; middle row, fixed background images; bottom row, fitted mass of each particle at sequential measurements.

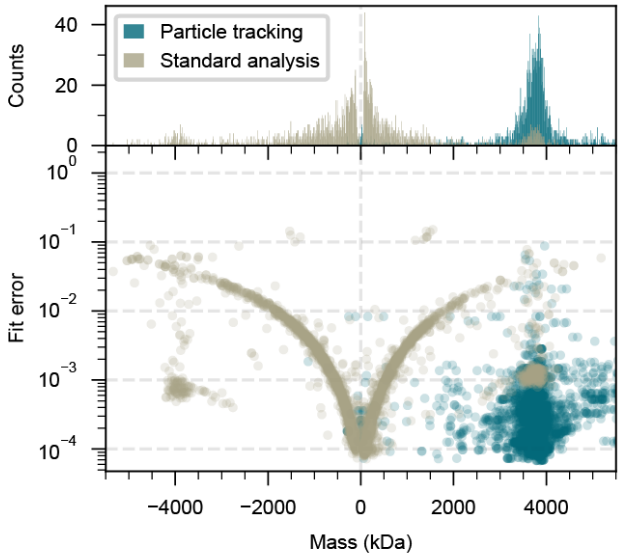


**Supplementary Fig. 7.** Particle tracking resolves the capsid mass distribution that standard analysis smears. Mass (top) and PSF fit error (bottom) for an empty AAV5 measurement (glass, pH 7.4) analysed by standard (ratiometric) analysis (grey) and by particle tracking (teal). Fit error is calculated as the sum of squared residual between the experimental signal and a mathematical PSF model.

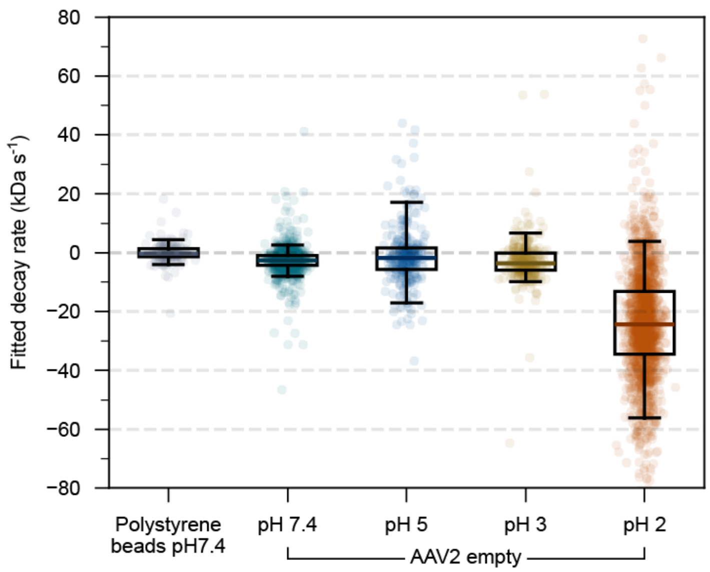


**Supplementary Fig. 8.** Rate of mass change of empty AAV2 capsids obtained from linear fits to individual mass traces in **Fig. 3c**, for polystyrene beads at pH 7.4 and empty AAV2 capsids on glass at the pH indicated. Each point represents the gradient from one fit to a capsids mass trace.

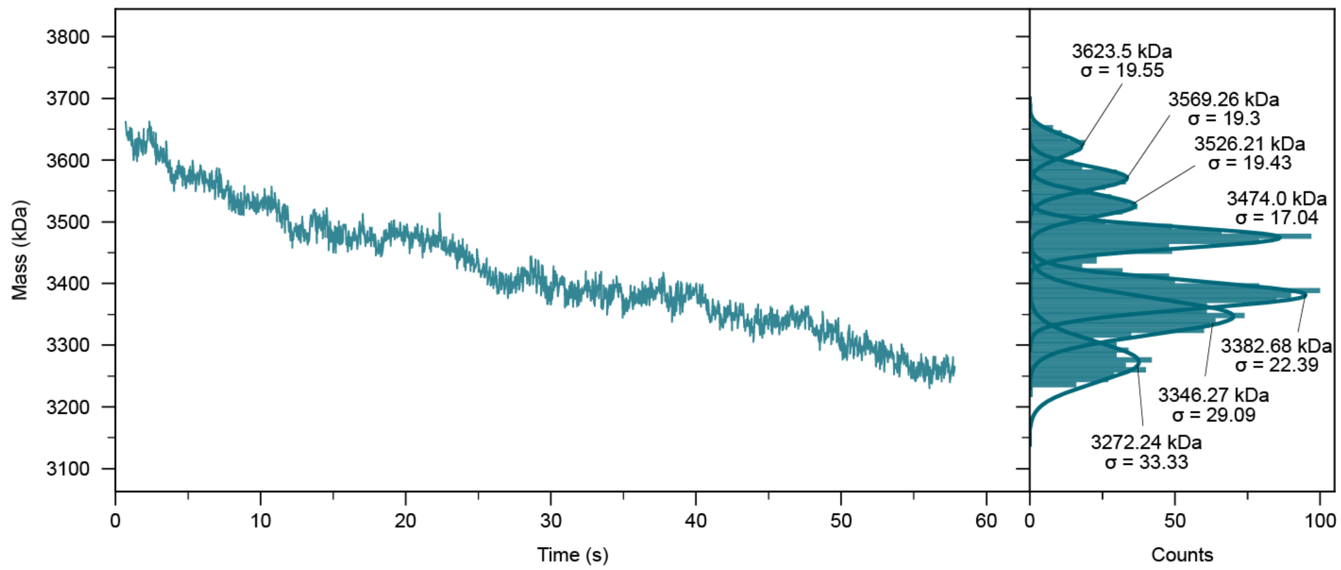


**Supplementary Fig. 9.** Mass over time trace of an empty AAV2 capsid (glass, pH 7.4) with Gaussian fits to peaks formed by long occupancy states in the mass distribution.

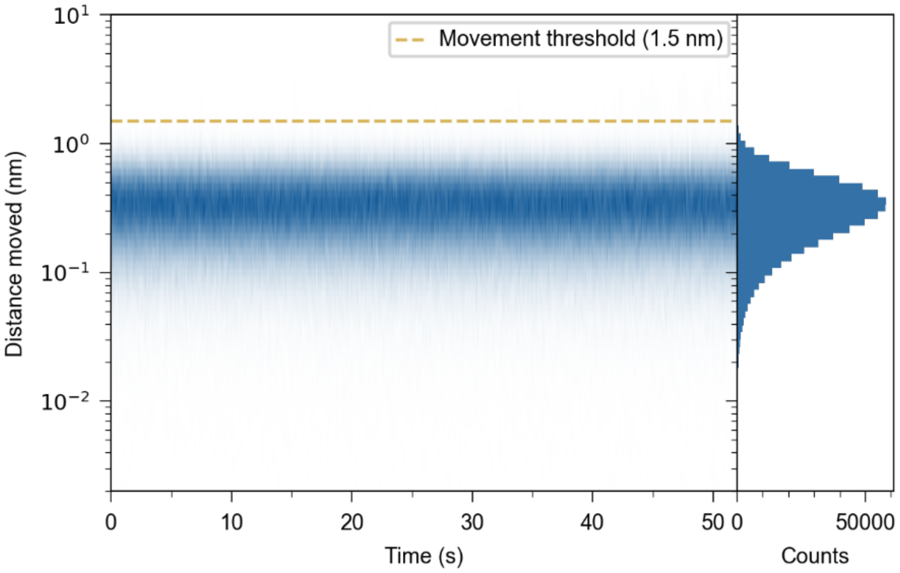


**Supplementary Fig. 10.** Frame-to-frame displacement of 262 stationary synthetic 3.5 MDa particles added to drift-corrected experimental buffer-only movies and analysed by particle tracking. The 1.5 nm movement threshold (gold dashed line) sets the used in **Fig. 3b** to classify real (significant) movements. The threshold is calibrated at 3.5 MDa; lower-mass particles have poorer localisation precision (**Fig. 1b**).

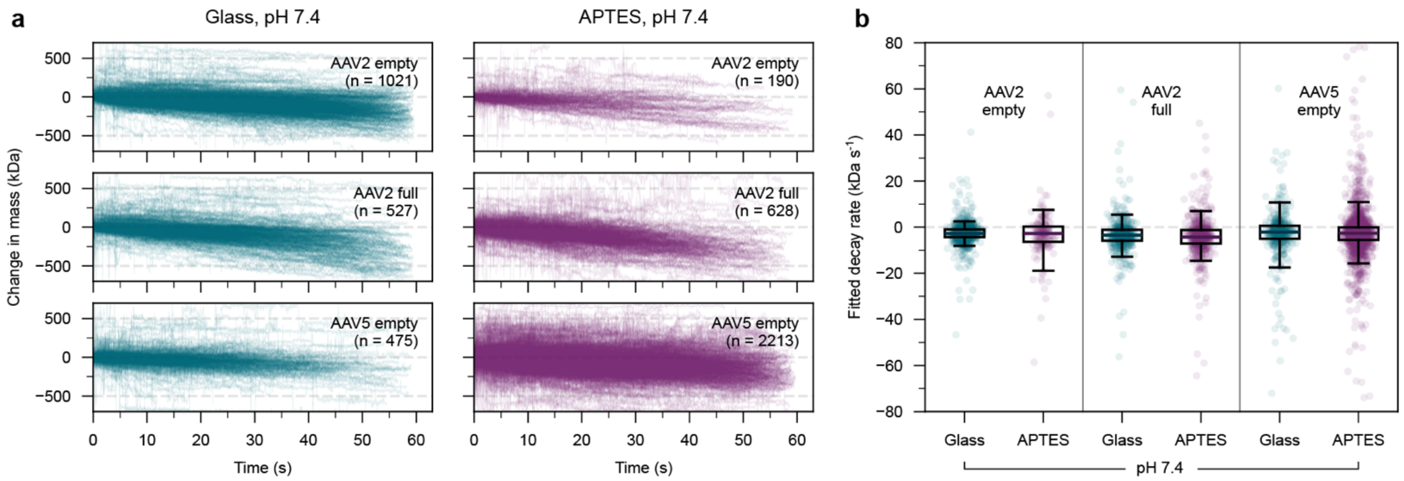


**Supplementary Fig. 11.** Mass change of AAV capsids on untreated and APTES-functionalised glass at pH 7.4. **a**, Change in mass relative to the first frame for empty AAV2, full AAV2 and empty AAV5 capsids on untreated glass (teal) and APTES-functionalised glass (purple) at pH 7.4. Each line represents the change in mass of one tracked capsid. **b**, Rate of mass change from linear fits to the traces in **a**, each point represents the gradient from one fit to a capsids mass trace.

| Instrument | Laser wavelength (nm) | $k_{\mathrm{off}}$ (s$^{-1}$) | ±95% Confidence interval (s$^{-1}$) |
|---|---|---|---|
| OneMP | 532 | 0.045 | 0.003 |
| TwoMP | 488 | 0.141 | 0.017 |
| Custom-built setup | 520 | 0.020 | 0.004 |
| Custom-built setup | 520 & 445 | 0.048 | 0.016 |

**Supplementary Table 1.** Dissociation rates of *D. spongiiphila* CS dimer products from hexameric precursors (with oxaloacetate) across different instruments (with different laser wavelengths).
